# Effect of antenatal tetramethylpyrazine on lung development and YAP expression in a rat model of experimental congenital diaphragmatic hernia

**DOI:** 10.1101/848556

**Authors:** Junzuo Liao, Wenying Liu, Libin Zhang, Qin Li, Fang Hou

## Abstract

Tetramethylpyrazine (TMP) is a chemical compound found in extracts derived from the Chinese medicinal plant. Due to its remarkable therapeutic effects, availability, and low cost and toxicity, TMP has been used to treat cardiovascular diseases and pulmonary hypertension in China. The aim of this study was to investigate the therapeutic effects and underlying mechanism of TMP on lung development using a rat model of nitrofen-induced congenital diaphragmatic hernia (CDH). Pregnant rats were divided into three groups: control, CDH, and CDH+TMP. Nitrofen was used to induce CDH. In the CDH and CDH+TMP, Fetuses only with left diaphragmatic hernias were chosen for analysis. Lung and body weight were recorded and lung histologic evaluations, image analysis, and western blot analysis of YAP, p-YAP and LATS1 were performed after lung processing. A marked abnormal structure was observed, as evidenced by pulmonary hypoplasia and vascular remodeling, in the CDH. These abnormalities were improved in the CDH+TMP. There were significant differences between the CDH and CDH+TMP in percentage of medial wall thickness, arteriole muscularization, radial alveolar counts, AA%, and alveolar septal thickness. YAP expression was markedly increased in the CDH compared to the control, which was not affected by antenatal TMP administration. However, prenatal TMP intervention significantly increased expression of LATS1 and phosphorylation of YAP in the CDH fetuses. Our results demonstrate that **a**ntenatal TMP administration improved vascular remodeling and promoted lung development in a rat model of CDH, potentially through increasing expression of LATS1 and phosphorylation of YAP.

## Introduction

Congenital diaphragmatic hernia (CDH) is an uncommon congenital malformation, occurring in 1 to 4 of every 10,000 pregnancies[1]. Although medical and surgical management of CDH has improved, CDH is still associated with a high mortality rate of 60 - 70% [2][2][4][5]. The disease name comes from the original abnormality involving a hole in the diaphragm, but over the last few decades clinicians have observed that the defect in the diaphragm is not a determinant of survival. The primary causes of mortality in CDH include pulmonary hypoplasia (PH) and severe persistent pulmonary hypertension of the newborn (PPHN) [6]. It was traditionally thought that PH was caused solely by herniation of the abdominal organs into the thorax through the pleuroperitoneal canals, which compresses the developing ipsilateral lung and limits the expansion of the contralateral lung. However, animal studies and evaluation of early human embryos have convincingly shown that abnormal pulmonary development in the embryonic phase is the primary defect [7].

Current postpartum care, surgery, or medical treatment strategies have not proven to be viable approaches for managing PH and improving the associated abnormal remodeling of the pulmonary vasculature. However, with the development of modern imaging technologies, CDH can now be accurately diagnosed in mid-gestation [8][9]. Therefore, it seems feasible to consider prenatal intervention in cases with poor prognosis. Fetal surgical interventions, such as fetoscopic temporary tracheal occlusion, are invasive, technically demanding, and limited by maternal and fetal risks [10]. Evidence has demonstrated that tracheal occlusion does not increase survival compared with standard postnatal care [11]. Therefore, less invasive approaches, such as antenatal pharmacologic treatment to stimulate lung growth and maturation, have been proposed and investigated in the laboratory [13][14][14]. Although some drugs have been found to improve pulmonary maturity and abnormal pulmonary vascular remodeling in animal models, the potential side effects of antenatal treatments, such as glucocorticoids [15] and sildenafil [16], limit their use. Therefore, it is necessary to explore other pharmacologic antenatal interventions to determine which have fewer side effects.

Tetramethylpyrazine (TMP, also called ligustrazine) is a chemical compound found in extracts derived from the Chinese medicinal plant Ligusticum wallichii (*family Apiaceae*). TMP possesses typical characteristics as a calcium antagonist. Due to its remarkable therapeutic effects, availability, and low cost and toxicity, TMP has been considered an effective therapy for various diseases, such as cerebral ischemia, cardiovascular diseases, and pulmonary hypertension in China [17][18]. Moreover, in recent years, TMP has been found to be an effective and safe treatment for fetal growth restriction (FGR) [19].

Most of our current understanding about the structural and molecular changes in CDH originated from experimental animal models[20]. Administration of the herbicide nitrofen (2,4-dichloro-phenyl-pnitrophenyl ether) to pregnant rats on embryonic day 9.5 (E 9.5) has been shown to result in PH and diaphragmatic defects in the offspring, both remarkably similar to human CDH[21][22].

The purpose of this study was to evaluate the effect of TMP on improving CDH-induced abnormal pulmonary vascular remodeling and PH in the nitrofen-induced CDH model. We also explored the possible underlying mechanism of TMP’s effects by measuring expression and activation of Yes-associated protein (YAP), which is an important protein in pulmonary development and vascular reconstruction.

## Materials and methods

### Experimental design and animal model

All animals were provided by the Institute of Laboratory Animais of Sichuan Academy of Medical Sciences and Sichuan Provincial People’s Hospital (Chengdu, China). The protocol was approved by the Committee on the Ethics of Animal Experiments of Sichuan Academy of Medical Sciences and Sichuan Provincial People’s Hospital (Protocol Number: 2018-198). All surgery was performed under sodium pentobarbital anesthesia, and all efforts were made to minimize suffering. This experiment was supported by Lilai Biotechnology (Chengdu, China). Twenty adult female Sprague-Dawley rats weighing 240 - 305 g (average, 283 g) were used. All rats were bred after a night of controlled mating. A sperm-positive vaginal smear confirmed mating and represented embryonic day (E) 0.5. CDH was induced in pregnant rats at E9.5 via intragastric administration of a single oral dose of nitrofen (125 mg; 99% purity; Zhejiang Chemicals, Ningbo, Zhejiang, China) that was dissolved in 2 ml of olive oil. Control rats received an equal amount of olive oil only. On E11.5, nitrofen-fed pregnant rats were randomly divided into two groups: CDH or CDH+TMP. TMP (80 mg/kg, Livzon Pharmaceutical Group Inc., China) was intragastrically administered to pregnant rats on E11.5 once a day, for 10 days. In total, our study included three groups of pregnant rats as follows: control (n = 5), CDH (nitrofen-induced CDH, n = 5), and CDH+TMP (nitrofen-induced CDH with antenatal TMP treatment, n = 5). Rat fetuses were delivered via Cesarean on E21.5 (prior to full term, E22) and immediately beheaded after being weighed. Under an anatomic stereoscopic microscope, fetal lungs were removed and the bilateral diaphragms were carefully examined for CDH. Lung tissue weight (LW) and body weight (BW) of each fetus were recorded. The left lungs of the fetuses with left CDH only were removed and processed for further analysis.

### Lung preparation

Lungs were placed into 4% paraformaldehyde, fixed at 4°C for 48 hours, and then embedded in paraffin for histological analysis. Paraffin-embedded fetal lungs were transversely cut into 5 μm sections with a microtome. Sections were stained with hematoxylin and eosin (H&E) and an elastin histochemical stain. Lung samples to be used for western blot analysis were stored at −80°C.

### Morphological analyses

#### Pulmonary vascular morphometric analysis

Lung slices were deparaffinized and hydrated using conventional methods[23]. The sections were stained in Verhoeff’s solution for 1 hour until the tissue sections were completely stained black. Then, the sections were washed in running tap water three times and soaked in 2% ferric chloride for 1 - 2 minutes, rinsed briefly with tap water, and checked for black elastic fiber staining and gray background under a light microscope. The slides were treated with 3% sodium thiosulfate for 5 minutes and rinsed in running tap water for 5 minutes. Slides were then counterstained in Van Gieson’s solution for 5 minutes, dehydrated, and rinsed in graded alcohols and xylene, cover slipped, and observed under a light microscope.

External diameter (ED) and medial wall thickness (MT) of small pulmonary arteries with a diameter of 20 ∼ 60 μm that were associated with terminal bronchioles and distal airspaces were quantified using Image-Pro Plus 6.0 (Media Cybernetics, Inc., Chengdu, China). ED was defined as the distance between the external elastic laminae, and MT was defined as the distance between the internal and external elastic laminae. Percentage of medial wall thickness (%MT) was calculated using the following formula: 2 × MT/ED × 100 [24][25]. Vessels with a measurable medial wall were considered muscularized; vessels without a medial wall were considered non-muscularized; and vessels with an incomplete medial wall were considered partially muscularized[15].

#### Lung maturation measurement

Lung growth was determined using the LW/BW ratio. The following morphological parameters were measured: (1) Radial alveolar count (RAC) was used as an index of alveolar proliferation and architectural maturity. RAC has been used to measure development of the terminal respiratory unit, as originally described by Emery and Mithal [25]. RAC was determined by counting the number of airspaces along a line drawn perpendicularly from the center of a terminal or respiratory bronchiole to the closest edge of the acinus (pleural or lobular connective tissue septum). (2) Percentage of lung alveolar area per unit area (%AA) was measured by image analysis using Image Pro Plus version 6.0 (Media Cybernetics, Inc., Chengdu, China). (3) Alveolar septal thickness was also measured.

### Western blot analysis

Lung tissues were homogenized in RIPA buffer supplemented with Complete Protease Inhibitor Cocktail tablets (Roche) and phosSTOP Phosphatase Inhibitor Cocktail tablets (Roche). Protein concentrations were determined using the Pierce BCA assay (Rockford, IL). Total protein (50 μg) was linearized in Laemmli sample buffer (Bio-Rad, USA) and then separated by gel electrophoresis using prefabricated 10% SDS polyacrylamide gels (Invitrogen). Proteins were then transferred to PVDF membranes (Hybond, USA). Immediately after transfer, the membranes were blocked with 5% bovine serum albumin (BSA) for 2 hours before antibody detection. Primary antibodies against YAP (1:500, CST), large tumor suppressor kinase 1 (LATS1) (1:300, Proteintech), phosphorylated (p)-YAP (1:1,000, CST), and β-actin (1:5000, Abcam) were incubated overnight at 4°C. The membranes were further incubated with a goat anti-rabbit secondary antibody (1:5,000, Abcam) at room temperature for 2∼3 hours followed by extensive washing. An enhanced chemiluminescence (ECL) kit (Thermo, USA) was used for antibody detection. All antibodies used in this study were diluted in phosphate buffered saline (PBS). The gel image analysis system (Tanon, China) was used for scanning analysis and the results are presented as relative expression of the target protein calculated as: target protein expression = integrated optical density value of target protein / internal reference integrated optical density value.

### Statistical analysis

Values are presented as mean ± standard deviation (SD). All data were statistically analyzed using SPSS, version 21 (SPSS, Inc, Chicago, III). Statistical analysis was performed using one-way ANOVA and the Chi-square test. Values of *P* < .05 are considered statistically significant.

## Results

### Incidence of CDH and pulmonary vascular remodeling

We determined the incidence of CDH in the three groups: none of the 80 fetuses in the control group presented with CDH; 48 out of 70 fetuses (68.6%) presented with CDH in the CDH group; and 51 out of 76 (67.1%) fetuses presented with CDH in the CDH+TMP group. There were no significant differences in CDH incidence between the CDH and CDH+TMP groups (*P* = .85). In the experimental groups, fetuses with left CDH only were included in our analyses (Table 1). Accordingly, the fetuses were further divided into three groups: controls (n = 80), CDH (n = 35), and CDH+TMP (n = 41).

**Table 1.** Pulmonary vascular morphometry.

| Group | N | ED (μm) | %MT | Wall structure (%) | | |
| --- | --- | --- | --- | --- | --- | --- |
|  |  |  |  | M | PM | NM |
| Control | 80 | 25.22±8.82& | 17.86±4.18 | 49.56±9.76 | 4.27±1.85 | 46.17±8.97 |
| CDH | 35 | 30.19±11.35 | 34.84±5.91* | 58.46±11.2<br>1 | 15.69±4.77 | 25.85±9.39§ |
| CDH+TMP | 41 | 27.27±9.87&& | 18.54±4.11** | 51.07±8.90 | 7.69±2.05 | 41.24±10.11§§ |

Values are expressed as $\bar{x} \pm s$ . M: fully muscularized; PM: partially muscularized; NM: non-muscularized. & $P = .57$ , && $P = .71$ , vs Control; \* $P < .01$ , vs Control; \*\* $P < .01$ , vs CDH; § $P < 0.01$ , vs Control; §§ $P < .01$ , vs CDH.

Comparison of pulmonary vascular morphometry showed that ED was not statistically different between the CDH group and the other groups (*P* = .57, P=.71). Fetuses in the CDH group had a significantly increased %MT and decreased non-muscularized vessels compared to the controls (*P* < .01, *P* < .01), whereas the fetuses in the CDH+TMP group showed significantly reduced %MT and increased non-muscularized vessels compared to the CDH group (*P* < .01, *P* < .01) (Table 1, Fig 1).

**Fig 1.**
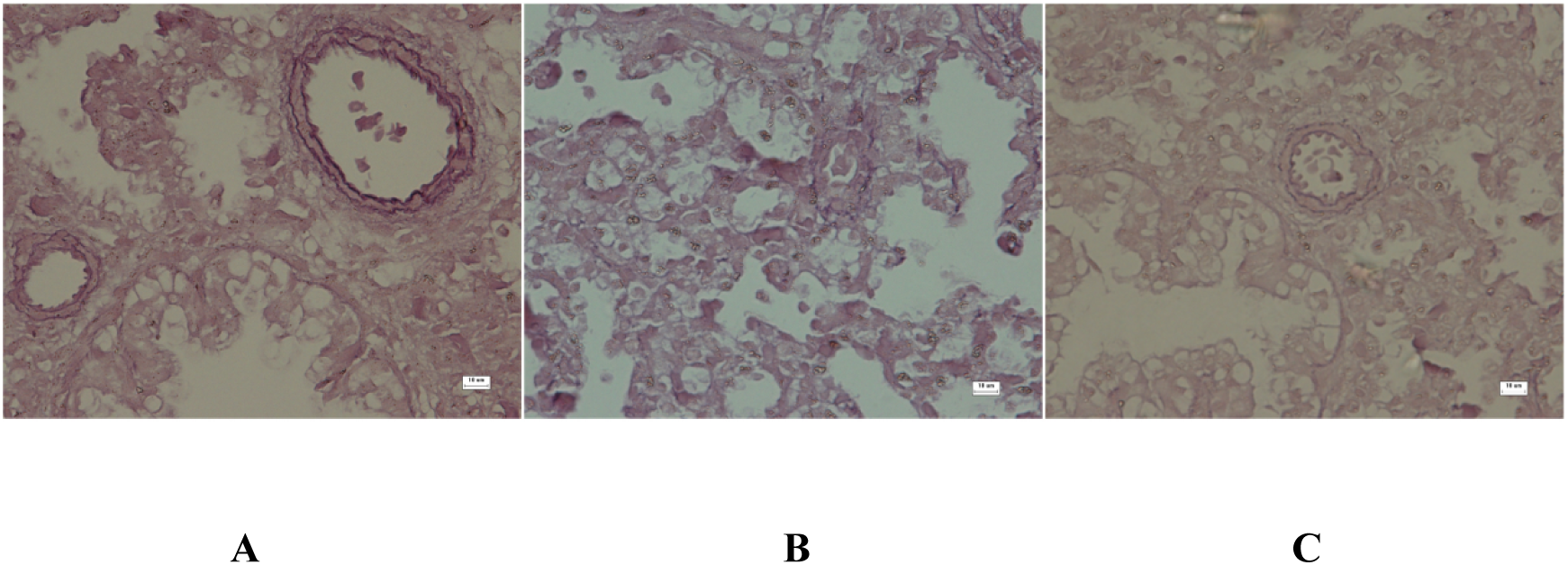
Medial wall thickness of the pulmonary artery was significantly increased in the CDH group (B) compared to the control group (A). Medial wall thickness was decreased in the CDH+TMP group (C) compared to the CDH group (B) (VVG, original magnification ×400).

### LW/BW ratio and lung morphometric analysis

Similar to our previous study, fetal lungs in the CDH group were markedly hypoplastic, as evidenced by lower LW/BW ratios, decreased RAC and %AA, and thicker alveolar septum compared to the CDH group. Treatment with TMP significantly promoted fetal lung development, but the development still lagged compared to controls. The results are shown in Table 2 and Fig 2.

**Table 2.**
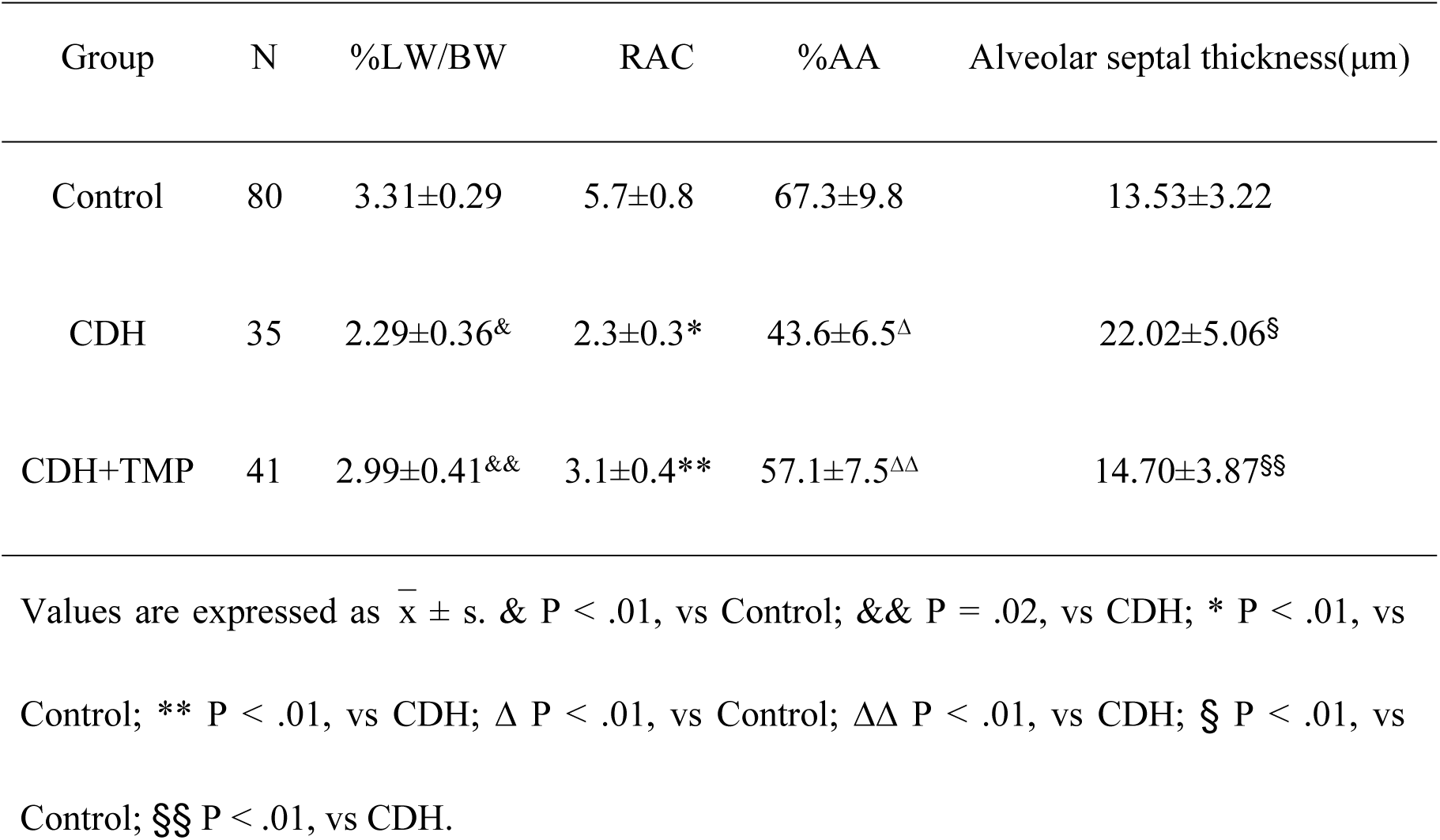
LW/BW ratio and lung morphometric analysis.

**Fig 2.**
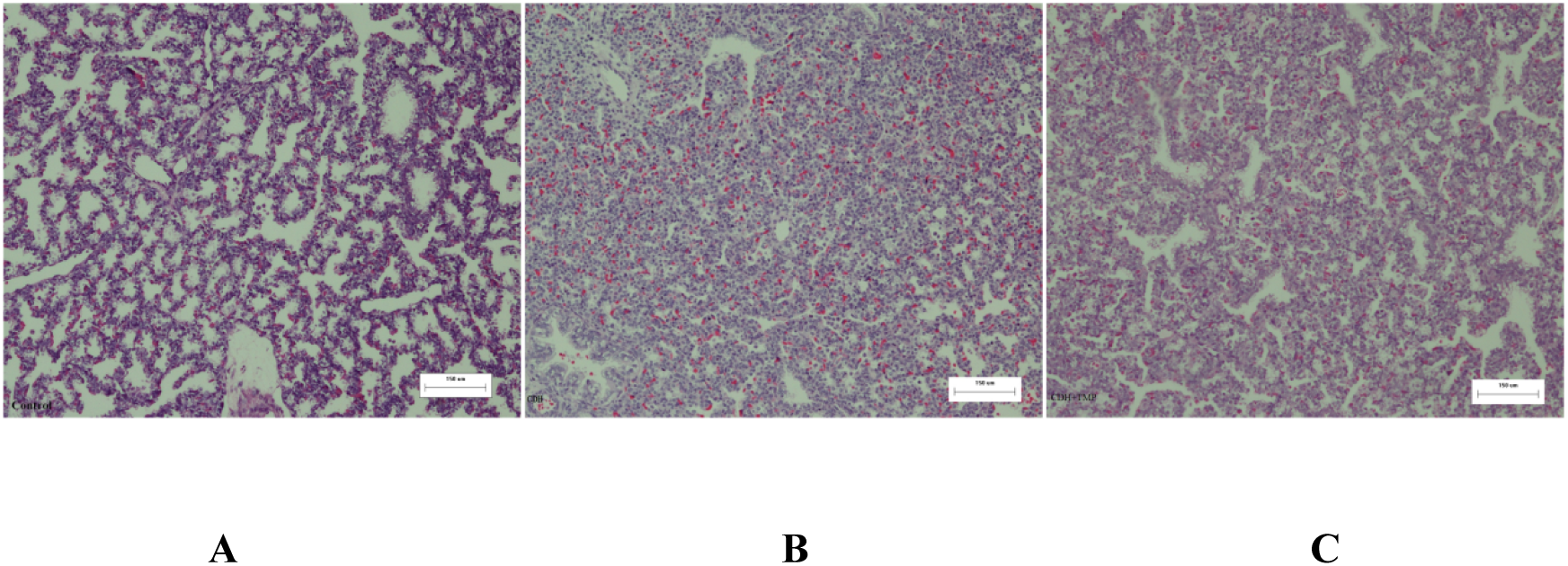
Lungs from the CDH group (B) were characteristic of the fetal canalicular stage, showing poorly formed saccules and thickened septal walls compared to lungs in the control group (A), which showed well-differentiated saccules and thin septal walls. Striking changes, including an increase in air saccule size, thin septal walls, and maturation of the pulmonary interstitium, (H&E, original magnification ×100) were seen in the CDH+TMP (C) groups.

### Western blot analysis of YAP, LATS1, and p-YAP

YAP expression was significantly increased in fetal lungs from the CDH group compared to the control group (*P* < .01), while there was no significant difference in LATS1 between the two groups (*P* = .65). TMP prenatal intervention did not significantly affect YAP expression (*P =* .28), but significantly increased LATS1 (*P* < .01) and p-YAP (*P* < .01) expression in the CDH lung tissues. Equal loading of electrophoresis gels was confirmed by β-actin staining of the stripped membranes (**Fig 3**).

**Fig 3.**
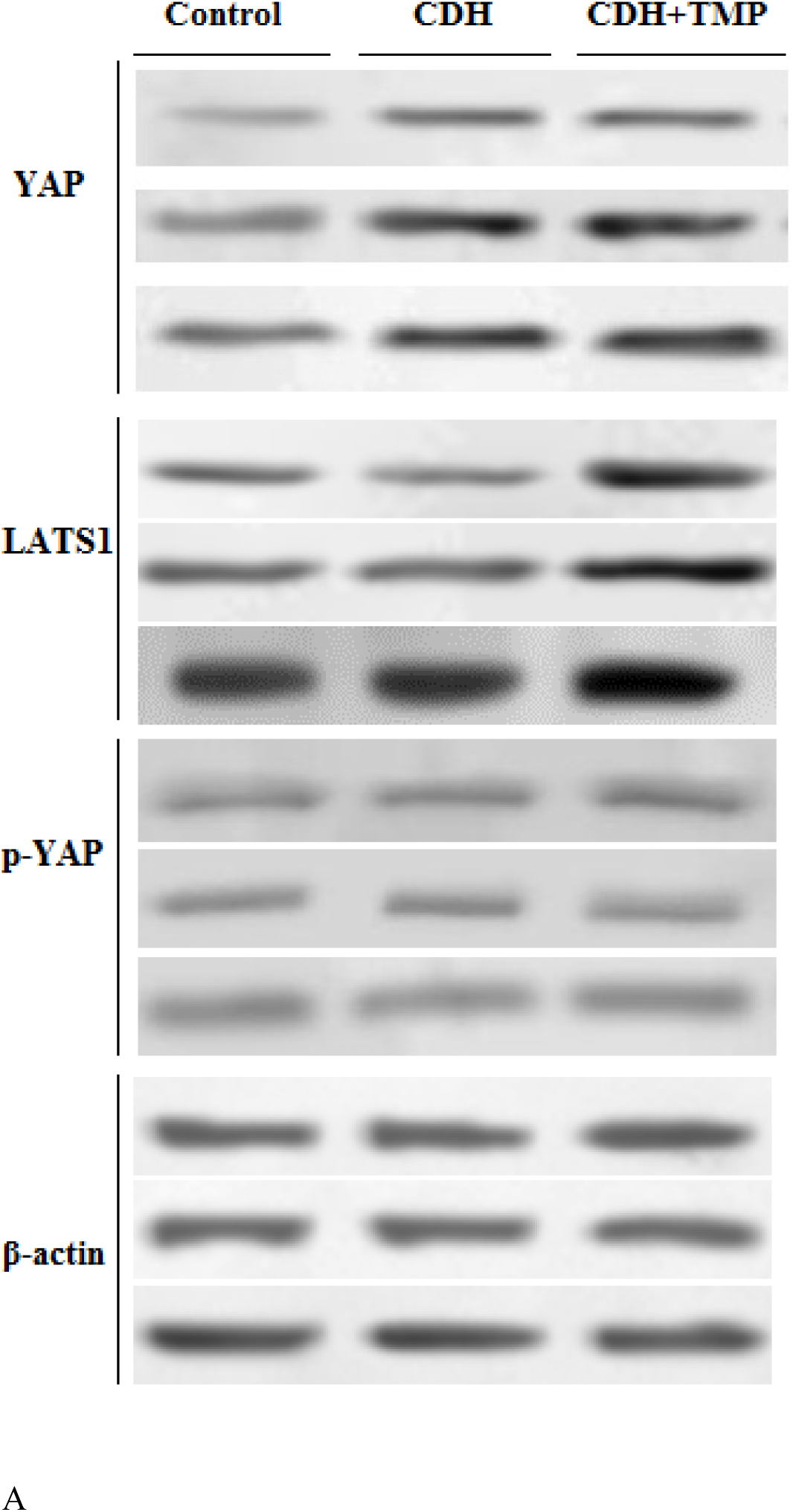
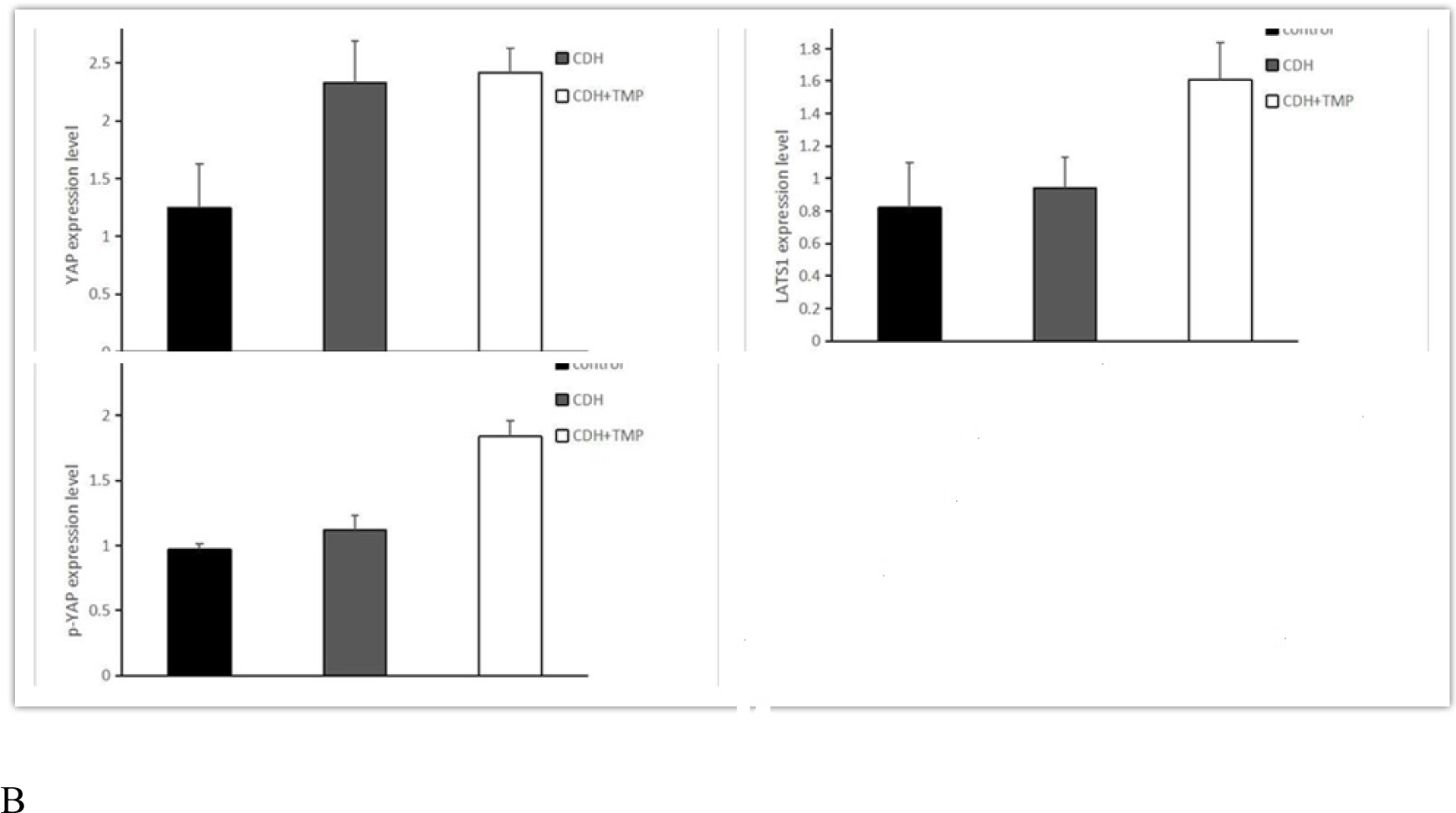
(A) Western blot analysis of lysates derived from control, CDH, and CDH+TMP lung tissues. (B) YAP expression was significantly increased in the lungs of the CDH group compared to the control group (P < .01). TMP prenatal intervention did not significantly affect YAP expression in CDH fetal lung tissue (P = .28), but significantly increased LATS1 (P < .01) and p-YAP (P < .01) expression.

## Discussion

TMP has been used in traditional Chinese medicine for many years to treat various diseases, including pulmonary hypertension, cardiovascular and neurovascular disease, FGR, and others. Therefore, we hypothesized that TMP could also be used to treat CDH with PH and PPHN. In this study, we used a rat nitrofen-induced CDH model to evaluate the effects of prenatal TMP administration on improving pulmonary vascularization. Our results indicate that prenatal TMP therapy significantly reduced medial thickness of small arteries and increased the number of non-muscularized arteries, while decreasing the number of fully or partially muscularized arteries in CDH rats. These data indicate that TMP decreases vascular remodeling, resulting in increased pulmonary blood flow, and further suggests that pulmonary hypertension in CDH rats can be alleviated by prenatal TMP therapy. However, the mechanism by which TMP inhibits pulmonary vascular remodeling in the CDH rat model remains unclear. Emerging evidence supports that YAP plays an important role in vascular remodeling and related cardiovascular diseases [27]. Therefore, we hypothesized that TMP alters YAP expression and activation in CDH.

In mammals, YAP is the key functional effector of the hippo pathway, which mainly comprises mammalian STE20-like protein kinase 1/2 (MST1/2), Salvador family WW domain containing 1 (SAV1), large tumor suppressor 1/2 (LATS1/2), Mps one binder (MOB1), YAP/transcriptional coactivator with PDZ-binding motif (TAZ), and transcriptional enhancer associate domain family members 1-4 (TEAD1-4) [28][29]. When the Hippo pathway is activated, the YAP/TAZ complex is phosphorylated by LATS1/2, which results in its nuclear exclusion, ubiquitination, and subsequent proteolytic degradation [30]. Hippo/YAP signaling plays an important role in cardiovascular development and vascular homeostasis [31]. Moreover, Hippo/YAP signaling has been found to contribute to vascular remodeling and related cardiovascular diseases, including pulmonary hypertension, atherosclerosis, aortic aneurysms, restenosis, and angiogenesis [27]. New evidence suggests that YAP regulates proliferation and survival of pulmonary arterial vascular smooth muscle cells (VSMCs) and pulmonary vascular remodeling [32][33]. In addition, LATS1 was found to be inactivated in small remodeled pulmonary arteries, as well as distal pulmonary arterial VSMCs in idiopathic pulmonary hypertension [32]. In our study, we found that upregulated YAP expression in the CDH rats was associated with increased pulmonary vascular resistance and altered pulmonary arterial muscularization. We also found that TMP treatment increased LATS1 expression and YAP phosphorylation. Therefore, we speculate that pulmonary vessel remodeling and pulmonary hypertension in CDH is partly due to an increase in LATS1 and YAP expression and activity.

YAP transcriptional targets often include positive regulators of cell proliferation and negative regulators of cell death. Thus, inactivation of Hippo signaling leads to organ enlargement, which is a signature phenotype of Hippo pathway activation [34][35]. However, in this study, we found that increased YAP expression in CDH lung tissues did not lead to increased lung size; rather, upregulation of YAP led to a decrease in lung size and an apparent cessation in development. A recent study suggested that early inactivation of the Hippo pathway during early stages of lung development resulted in a sharp decrease, rather than the expected increase, in lung size [36]. Furthermore, researchers found that, despite nuclear YAP localization in the epithelium, Shhcre;Lats mutant rats had smaller lungs with halted development after primary branch formation, one of the more severe lung developmental phenotypes. This developmental defect is likely attributed to disrupted localization of apical-basal polarity determinants, cell adhesion molecules, extracellular matrix components, and spindle misorientation in dividing cells. Instead of a single cell layer of epithelium that is critical for effective extension and growth of the branches, Shhcre;Lats mutant rats showed a multilayered epithelium with cells protruding into the lumen. Similar phenotypes were also described in the kidneys and salivary glands of transgenic animals over-expressing YAP or mutants with a LATS1/2 deletion [37][38]. These findings suggest that in branching organs, such as the lung, kidney, and salivary gland, there is a primary role for Hippo signaling to maintain an organized epithelium, which is critical for degerming organ size [36]. Moreover, increased Yap activity could lead to impaired differentiation and maturation of lung epithelial cells and decreased surfactant proteins [39][40], all of which are in accordance with many disease manifestations in CDH lung tissues.

Interestingly, we found that antenatal administration of TMP was beneficial for improving PH, as evidenced by the LW/BW ratio, alveolar septal thickness, RAC, and %AA in the CDH+TMP group compared to the CDH group. We speculate that these results are related to increased LATS1 expression and inhibition of YAP activity.

This study is the first to report the effects of prenatal TMP administration on lung development in a rat model of CDH. We revealed a significant role of Hippo signaling in CDH-associated PH and pulmonary hypertension. We also found that antenatal nitrofen exposure increased YAP expression and structural abnormalities in the lung, including abnormal vascular remodeling and impaired alveolarization. We also noted that antenatal TMP treatment promoted lung development and improved vascular remodeling. Although further studies are needed to determine the exact mechanisms of CDH-induced PH and antenatal TMP administration in improving lung structure, the current findings suggest that increased YAP activity is associated with delayed pulmonary development and abnormal vascular remodeling, and antenatal TMP therapy improves lung structure and function via increasing LATS1 expression and phosphorylation of YAP.

There are several limitations to note in our study. There was a lack of data regarding the whole process of fetal lung development, and therefore we could dynamically reflect the changes of the Hippo signaling pathway in lung development. Furthermore, there was a lack of direct evidence supporting the relationship between the Hippo signaling pathway, CDH-induced PH, and TMP prenatal intervention. These limitations need to be addressed in future studies.

## Availability of data and materials

The authors declare that all data supporting the findings of this study are available within the article or from the corresponding authors on reasonable request.

## Acknowledgements

We wish to thank Honghui Jia and Lin Wang for kindly guiding the establishment of animal models. We thank Lin Jiang for statistical analysis.

## Authors’ contributions

JZ Liao is first author. WY Liu obtained funding. JZ Liao, Q Li, LB Zhang, and WY Liu designed the study. JZ Liao, Q Li, and LB Zhang collected the data. JZ Liao and Q Li were involved in data cleaning and verification. JZ Liao and Q Li analyzed the data. JZ Liao drafted the manuscript. WY Liu, JZ Liao, and F Hou contributed to the interpretation of the results and critical revision of the manuscript for important intellectual content and approved the final version of the manuscript. All authors have read and approved the final manuscript. JZ Liao and WY Liu are the study guarantors.

